# Human color constancy in cast shadows

**DOI:** 10.1101/2024.02.10.579744

**Authors:** Takuma Morimoto, Masayuki Sato, Shoji Sunaga, Keiji Uchikawa

## Abstract

Most natural scenes contain cast shadows to a varying extent. Illuminant conditions inside and outside the shadow typically differ largely both in intensity and in chromaticity. Nevertheless, our daily experiences suggest that colored materials appear to have the same color in shadows even though the reflected light from the material might be very different. Two experiments were conducted to reveal mechanisms that support human color constancy in cast shadows. In all experiments we built a real scene that consisted of colored hexagons illuminated by two independent liquid crystal projectors simulating “sunlight” and “skylight”, respectively. A part of the scene included a cast shadow under which observers were instructed to change the luminance and the chromaticity of a test field so that it appeared as a full-white paper under the shadow. The color of the skylight was manipulated, testing if our visual system uses a prior that the skylight is typically bluish or yellowish to achieve color constancy. We also created a condition where a cast shadow is not recognized as a shadow. Results showed that color constancy generally holds well in shadows and changing skylight color had little effect. Recognizing a cast shadow as a shadow also had no effect. Overall, these results are consistent with our daily experiences that we stably judge objects’ color even in shadows, providing a key step to reveal mechanisms of color perception in real-world scenes where lighting conditions spatially vary.

## 1. Introduction

Our ability to judge the surface color of an object is stable despite large variations of lighting environments in the real world. This is widely known as color constancy. The extent to which color constancy holds has been measured under diverse experimental conditions, and various strategies to “undo” the influence of illumination have been proposed (Foster, 2001). However, one significant limitation in those studies is that scenes were lit by a single light source (e.g. Maloney & Wandell 1986; Morimoto et al. 2016; Morimoto et al. 2021), and there are only a handful of behavioral studies investigating mechanisms of color constancy when there are multiple illuminations in the scene (Yang & Shevell 2003; Smithson & Zaidi, 2004; Boyaci, Doerschner, & Maloney, 2004; Doerschner, Boyaci & Maloney, 2004; Lee & Smithson 2012).

Yet, such multi-illuminant conditions are not rare in natural environments. For example, sunny outdoor scenes typically contain two major illuminants, a sunlight and a skylight (Preetham, Shirley, & Smits 1999; Tian et al., 2016; Wilkie et al, 2021). Color constancy in a multi-illuminant environment is inherently more challenging for our visual system than in a single-illuminant environment because the influence of illuminant needs to be inferred differently for different spatial locations. One further complication is that the occlusion of one light generates a cast shadow under which the influence of the other light becomes dominant, creating a complex spatial lighting variation. Figure 1 shows a photo taken by an author at Ookayama Campus of the Tokyo Institute of Technology in Japan. Readers can presumably recognize that the pedestrian crossings are white regardless of whether they are under the sun (upper part) or under the shadow (lower part). However, as depicted by squares in the figure, the pixel value for the region in the shadow is dark blue because it exclusively reflects the skylight, which is drastically different from the pixel value of the region under the sun. This example effectively shows that our visual system can stably judge surface colors under such spatial variations of illuminations. Lightness, brightness and shape perception in shadows have been relatively well investigated (MacLeod, 1940; Adelson, 1999; Soranzo & Agostini, 2004; Knill et al., 1997; Mamassian et al., 1998). However, rather surprisingly, little empirical study is available for color constancy in shadows (Newhall et al., 1958) even though the idea was documented a few hundred years ago (Mollon, 2006).

**Figure 1:**
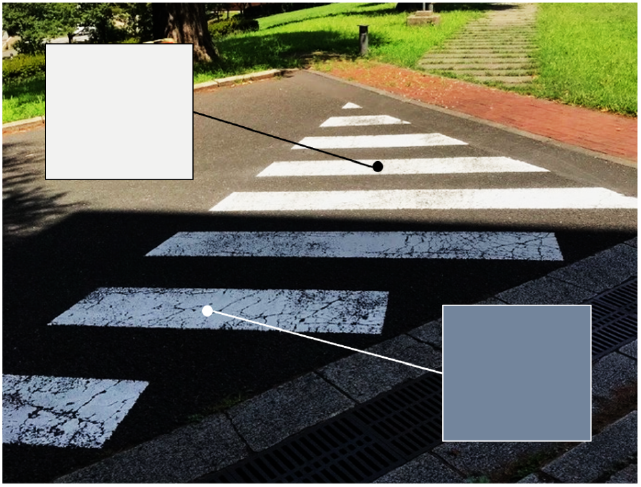
White pedestrian crossings in the picture would appear white to our eyes regardless of whether they are under the shadow or not. However, as shown in two squares, the pixel colors inside and outside of the shadow are drastically different. A picture taken by an author at Ookayama Campus of the Tokyo Institute of Technology.

To better understand the computational challenge our visual system faces in shadows, it is helpful to think about their physical characteristics. Measuring daylight spectra has been one active domain in the history of color vision studies (e.g. Judd et al. 1964; Hernández-Andrés et al. 2002), but there has been little research that specifically measured lights that reach cast shadows. To address this, we previously measured the spectral composition of lights reaching a cast shadow (i.e. skylight) in an outdoor environment and lights reaching regions under the direct sun (i.e. direct sun plus skylight) from dusk till dawn (Morimoto et al, 2022). As expected the measurements revealed that there are large spectral differences between two regions. On sunny days, skylights are the dominant light source that reach shadowed regions, and thus their spectra have high energy around a short wavelength region. In contrast, the non-shadowed region receives much brighter sunlight, and its chromaticity is located around white or slightly yellowish region in a color space. This complexity that shadows create naturally attracted researchers in the field of computer vision, and a wide range of methods for automatic detection and removal of shadows has been developed over decades (Lu & Drew, 2005; Wang et al. 2017; Qu et al., 2017; Cun et al. 2020; Liu et al., 2021; Le & Samaras, 2022), independent of more general algorithms and machine learning models to estimate multiple illuminants from a given scene (Gijsenij et al. 2012; Qu et al. 2015; Das et al. 2021; Wang et al. 2022). It is thus interesting to ask how humans effortlessly detect and discount the influence of a shadow.

The purpose of the present study was to understand the mechanisms to perceptually discount the influence of illumination in cast shadows. To answer this question, we conducted two psychophysical experiments. For both experiments, we constructed a real scene that was lit by two independent illuminants simulating “sunlight” and “skylight”, which were separately provided by two liquid crystal projectors. The scene contained a cast shadow under which an achromatic setting was measured. Overall, we asked three questions in this study.

1. How well does color constancy work in cast shadows? (Experiment 1)
2. Does the color of the skylight affect the degree of color constancy? (Experiment 1)
3. Does the recognition of a shadow have an effect? (Experiment 2)

Regarding the first point, due to the absence of recent empirical assessment, we felt that formally measuring color constancy in shadows should be a first step. The second point was to test the idea that human observers use statistical chromatic regularities in shadows in natural environments. This was directly inspired by a past empirical measurement showing that the color of skylight is dominantly bluish (Morimoto et al., 2022), but this is a wider interest to the field especially because in color constancy literature there has been an analogous argument regarding the daylight prior though its existence is still inconclusive (Delahunt, & Brainard, 2004; Pearce et al., 2014; Weiss, Witzel, & Gegenfurtner 2017; Morimoto et al., 2021a). The third point is based on our hypothesis that detection and recognition of the shadow would be required to notice that the shadowed region is illuminated differently from other regions in a scene. We achieved the condition by outlining the edge of the shadow (penumbra) with black surfaces, inspired by Hering’s outlined shadow (Hering, 1874/1964).

## 2. General Method

### 2.1 Experimental Apparatus

For all experiments, we set up a real scene with two independent liquid crystal projectors (HI-04, 1920 × 1080 pixels, 3600 lumens, DR. J Professional, Kent, Germany) which simulated “sunlight” and “skylight”. Overall there were two types of scenes. Figure 2A shows a set-up used in Experiment 1. The scene consisted of a sheet of colored hexagons arranged without a spatial gap. The central cup-shaped black object was placed to generate a cast shadow to the right part of the sheet. The sheet contained two test fields (holes) as shown inside the magenta square. Below the sheet, we placed an experimental monitor (ColorEdge CG2420, 24.1 inches, 1920×1200 pixels, 60 Hz, EIZO, Ishikawa, Japan) shown inside the blue square by which the chromaticity and the luminance of the test fields were modulated. The test fields extended approximately 2.5 degrees horizontally. We made sure that (i) from the observers’ viewpoint the holes were perceived as surfaces (rather than holes) and that (ii) it appeared to observers as if the color of the surface changed when the color of the monitor changed. In Experiment 1, both projectors were set on in one condition, and only the right projector (skylight) was set on in another condition. Physical properties of test surfaces and illuminants are described in the next subsection.

**Figure 2:**
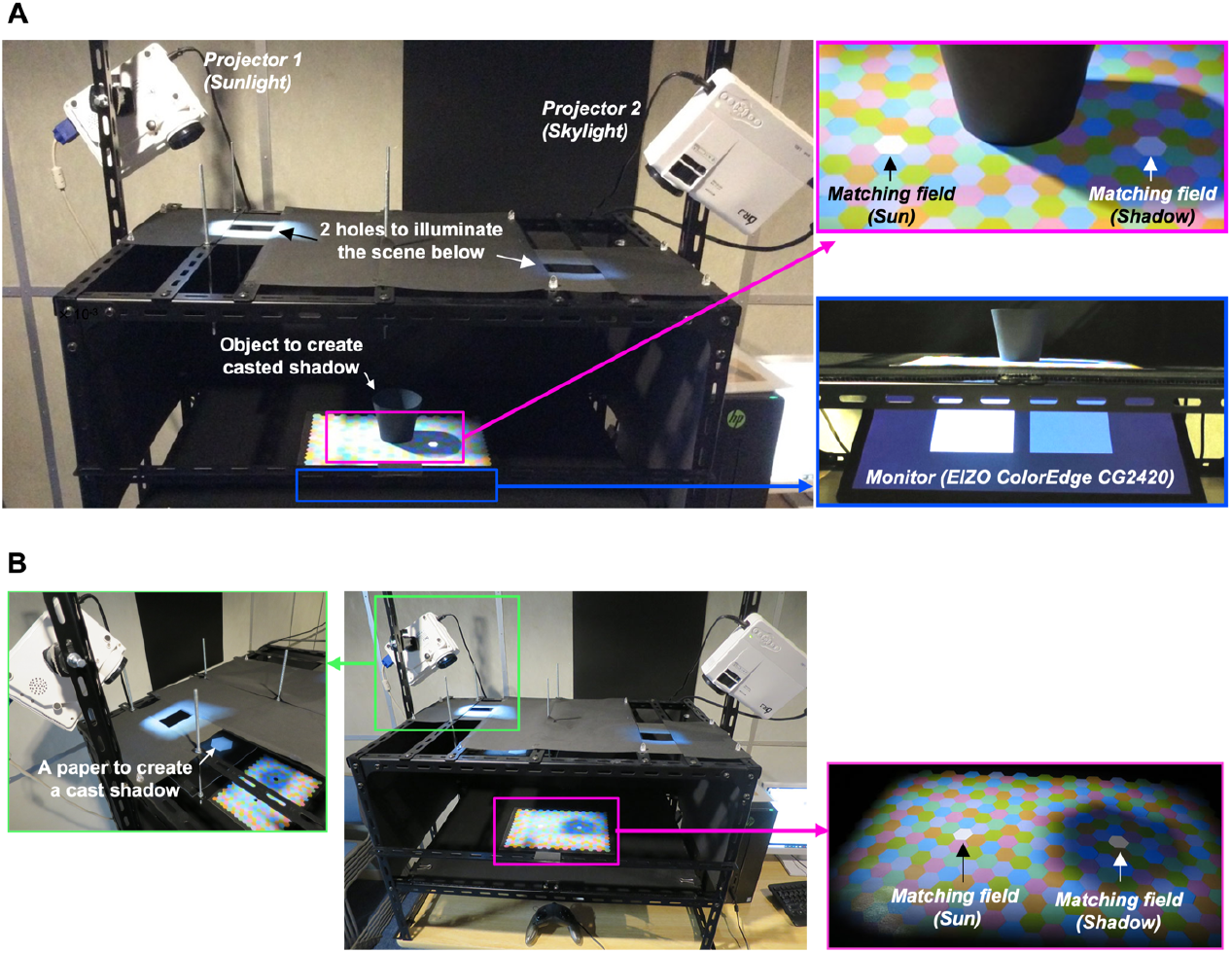
(A) Experimental setup for Experiment 1. Lights emitted from two projectors reach the scene. The black object placed at the center of the scene created a cast shadow. (B) Setup for Experiment 2. The center black object was removed, and a white paper was placed on an optical path between a projector and the scene.

Figure 2B depicts a set-up used in Experiment 2. We removed a center black object from the scene because the purpose of this experiment was to test whether color constancy holds even when a shadow is not recognized as a shadow (by outlining the shadow with black hexagons). If instead we had an object as in Figure 2A, observers would notice that there should be a cast shadow, and consequently shadowed regions are likely to remain perceived as shadows regardless of our manipulation. Thus to create a cast shadow on the sheet, we placed a white paper (seen in the green square, labeled as “a paper to create a cast shadow”) that blocked the sunlight emitted from a left projector. Otherwise the scene configuration was identical to Figure 2A. In Experiment 2, both left and right projectors were always set on.

Using a spectroradiometer (CS-2000, Konica Minolta, Tokyo, Japan) with 401 spectral channels (380 - 780 nm, 1 nm step), we performed spectral calibration to map RGB values to cone excitations and gamma correction to linearize monitor and projector outputs. The calibration was done separately for the experimental monitor and each of two LCD projectors. Stockman & Sharpe 2-degree cone fundamentals were used for the computation of cone excitations (Stockman & Sharpe, 1999; Stockman & Sharpe, 2000).

### 2.2 Properties of test surfaces and illuminants

The left and right projectors were used to simulate the sunlight and the skylight, respectively. The sunlight was fixed to a single spectral composition throughout the study (Figure 3A). The skylight was set to one of five spectra whose spectral compositions are shown in Figure 3B. These five colors for the skylight were chosen to represent typical (blue, yellow and white) and non-typical variations (green and magenta). The luminance of sunlight was 41.9 cd/m^2^ and skylights were all substantially darker than this, 3.12 cd/m^2^, 3.14 cd/m^2^, 3.15 cd/m^2^, 3.15 cd/m^2^, 3.00 cd/m^2^, for white, yellow, blue, magenta and green, respectively. These spectral distributions and luminances were measured by placing a BaSO_4_ white calibration plate to matching fields.

**Figure 3:**
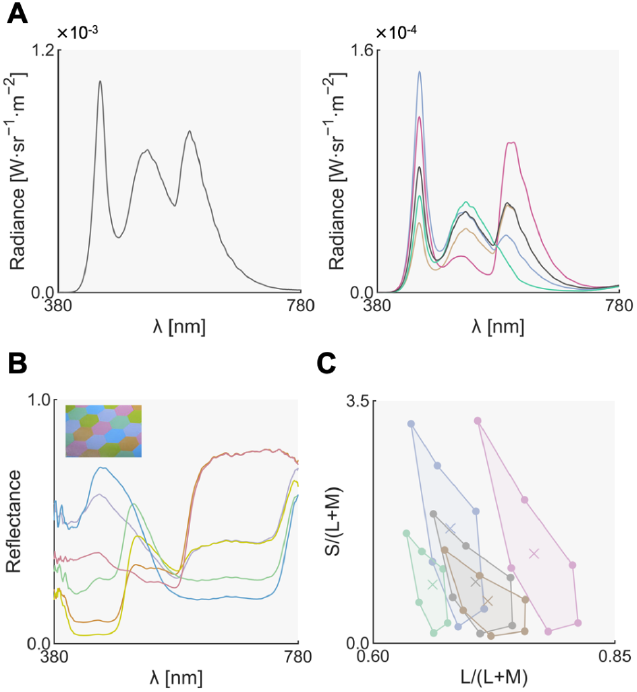
(A) Illuminant spectra of the sunlight (left panel) and skylights (right panel). Notice the difference in the y-axis range. (B) Spectral reflectances of six colored hexagons. (C) Filled circles show chromaticities of six colored hexagons under five skylights. The cross symbols show the chromaticities of five skylights.

We printed a sheet which contained six colored hexagons using a color laser printer (SPC840, RICOH, Tokyo, Japan) as shown in the upper-right corner of Figure 2A inside the magenta rectangle. We chose six colors so that they are visually distinct from each other when placed under the sunlight. Then, we measured the spectral reflectance of each color sample using a spectrophotometer (CS-2000, Konica Minolta, Tokyo, Japan) from 400 nm to 780 nm in 1 nm step (Figure 3B). Their MacLeod-Boynton chromaticities (MacLeod & Boynton, 1979) under five skylight illuminants are shown in Figure 3C. The vertices show chromaticities of six colors and the colors of the data points represent the color of the skylight. In this figure, the L/(L+M) and S/(L+M) of the equal energy white corresponds to 0.7078 and 1, respectively.

### 2.3 Observers

Three and twelve observers were recruited in Experiments 1 and 2, respectively. Two observers in Experiment 1 participated in Experiment 2, but otherwise no observers joined the other experiment. There was one female, and the rest were male observers. The mean and standard deviation of observers’ age were 57.0 and 8.72 in Experiment 1 and 28.3 and 14.8 in Experiment 2. All observers were screened to have normal color vision using Ishihara pseudoisochromatic plates and normal or corrected-to-normal visual acuity. Three observers in Experiment 1 and two observers in Experiment 2 were authors, and another observer in Experiment 2 were aware of the purpose of the experiment. Other observers were naive to the purpose of the experiment and had no or little experience in psychophysical experiments and no specialized knowledge about research in human color vision. In both experiments all observers completed all conditions. Informed consent was obtained from each observer before the start of the experiment. Observers were offered to take breaks during the experiments, and observers could stop the participation at any point.

### 2.4 Task and procedure

We used the method of achromatic adjustment in all experiments (Brainard, 1998). The original method required the adjustment of chromaticity only, but in our experiments, observers were asked to adjust the luminance and chromaticity of a test field until it appeared to be a full-white surface placed under a test illuminant (so-called “paper match” criterion; Arend & Reeves, 1986). Observers completed adjustments for both left and right test fields shown in Figure 2A, but in this paper we report only the matching results for the right test field as this was the main interest of the study. Therefore, throughout this paper, choosing a test chromaticity that matches the chromaticity of the skylight means a good color constancy.

For each illuminant condition, before starting the adjustment, the observer adapted to the scene for 60 seconds. For each trial the initial chromaticity and luminance of the test field were randomly selected. Observers were given no time limitation for the adjustment. The order of experimental conditions was randomized, but observers noticed when the illuminant color changed. More detailed experimental conditions are described in following experimental sections.

## 3. Experiment 1

### 3.1 Experimental condition

Figure 4 shows experimental scenes used in Experiment 1. All images were taken from approximately the observers’ viewpoints. Rows show the variation of skylight colors (white, yellow, blue, magenta, and green from the top). Panel A shows a “sunlight & skylight condition” where both left and right projectors were set on. Panel B shows a “skylight condition” where the scene was illuminated by the skylight only emitted from the right projector. This skylight condition served as a control condition in this study because the scene was lit by only a single illumination, resembling traditional color constancy experiments. Therefore, we took the degree of color constancy in this condition as a baseline constancy level. One block consisted of 5 skylight color conditions, and one session consisted of two blocks (sunlight & skylight, skylight-only). All observers completed 7 sessions in total.

**Figure 4:**
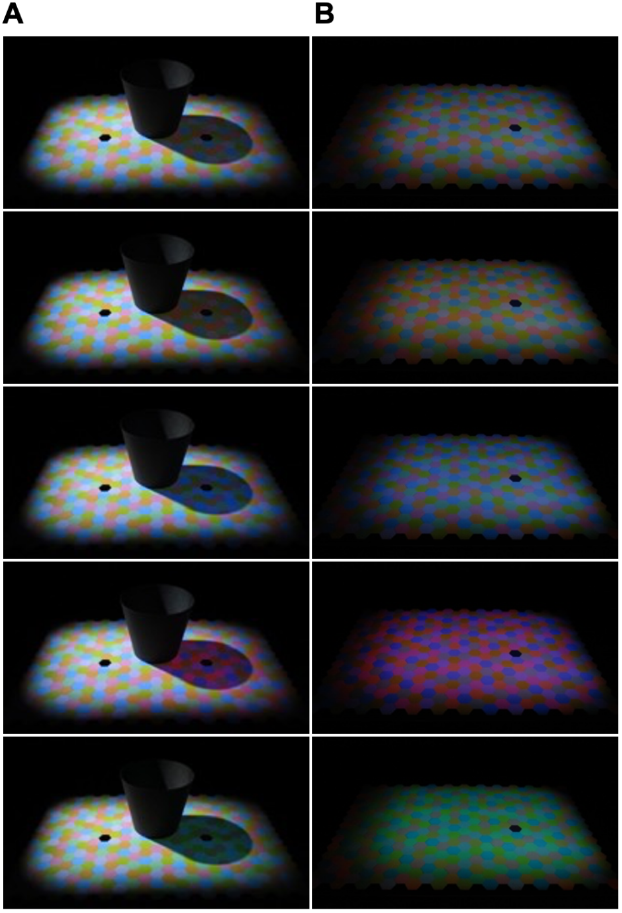
Experimental scenes used in Experiment 1. (A) Main condition (sunlight & skylight condition) where the scene was lit by two independent light sources. The black object was placed around the center of the scene to create the shadow. (B) Skylight condition (control) that is lit by the light emitted from the right projector only. This is a single-illuminant scene, in which a baseline color constancy level was measured.

### 3.2 Results

Figure 5 shows achromatic settings made by each observer (KU, MS and SS) for the sunlight & skylight condition (Panel A) and the skylight condition (Panel B). Circles are observer settings and crosses are chromaticities of the skylights. When the circle and the corresponding cross point match, that would indicate good color constancy. There was a general tendency that the observer settings shift towards the chromaticity of skylights, showing that color constancy worked to varying degrees. Note that these results are achromatic adjustments made for the right matching fields placed in the cast shadow (as shown by yellow arrows in this figure). We also analyzed the matching data for the left test files for the sunlight & skylight condition, but achromatic adjustments were unsurprisingly clustered around the chromaticity of sunlight regardless of the color of skylight because the skylight was much weaker than the sunlight and had a minimum effect on the left test field.

**Figure 5:**
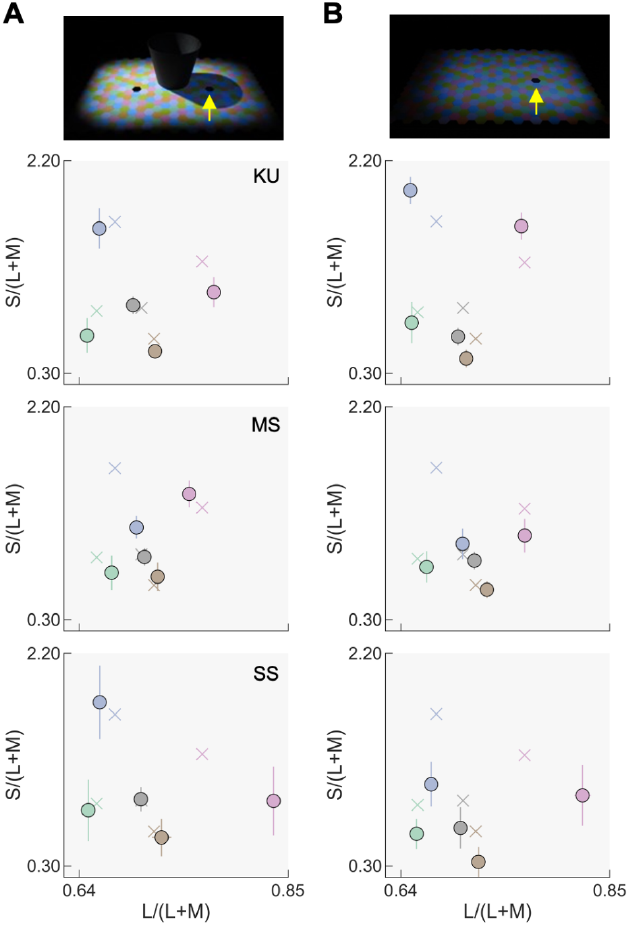
(A) Observer settings for three observers (KU, MS and SS) made for the right field for the sunlight & skylight condition. Each data point shows average across 7 settings. (B) Observer settings for skylight-only condition.

Next, as illustrated in Figure 6A, the degree of color constancy was quantified using the distance between chromaticity of the illuminants (defined as *b*) and the distance between chromaticity of the achromatic settings made by observers (defined as *a*). The reference illuminant was always the white skylight. To approximately equate the scale along horizontal and vertical axes, we divided each axis by the standard deviation of observer settings for the white skylight condition, separately for each observer. The constancy index *CI* was defined as equation (1). This definition is equivalent to a Brunswick ratio that incorporates the vector angle between perceptual illuminant shift and physical illuminant shift (Foster, 2011).

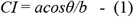

**Figure 6:**
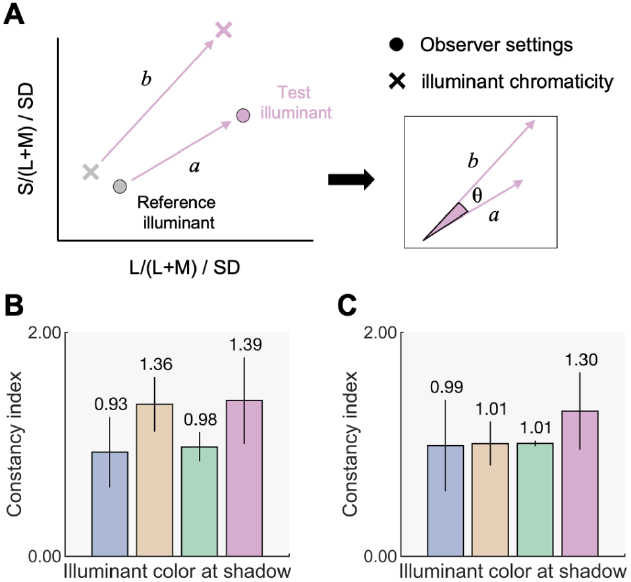
(A) How to compute a constancy index. Here, *b* denotes the distance between chromaticities of test illuminant (blue, yellow, green or magenta) and reference illuminant (white illuminant) and a denotes the distance of achromatic settings. Constancy index was defined as *acosθ/b*. (B) *CIs* for sunlight & skylight condition. The value was averaged across three participants and the error bars show ±1.0 S.E. (C) Mean *CIs* across three participants for the skylight condition. The error bars show ±1.0 S.E.

Constancy indices were first computed separately for each observer, and Figure 6B and 6C show indices averaged across all three observers for the sunlight & skylight condition and the skylight only condition, respectively. Overall, we observed a high degree of color constancy, but there are no obvious trends regarding the effect of skylight color and the number of light sources (sunlight & skylight or skylight only). To statistically test the effects we ran two-way repeated-measures analysis of variance (ANOVA) with skylight colors (blue, yellow, green or magenta) and illuminant condition (sunlight & skylight or skylight only) as the within-subjects factors for the *CIs*. The main effects of the color of skylights and illuminant condition were not significant (*F*(3, 6) = 0.96, *p* = 0.470, *F*(1, 2) = 0.72, *p* = 0.486, respectively). The interaction between the two factors was not significant, either (*F*(1, 2) = 0.56, *p* = 0.53).

This analysis concludes that human color constancy holds even in cast shadows. It is interesting that the degree of color constancy is as good as color constancy measured under the skylight only condition. There might be little cost for a visual system to deal with multiple illuminant conditions. The color of skylight had virtually no effect, which does not support the idea that humans use a prior about the color of illuminant reaching cast shadows in natural environments.

## 4. Experiment 2

The primary purpose of Experiment 2 was to test whether the recognition of the presence of a shadow provides a cue to know that the cast shadow is illuminated differently from other regions of the scene. To test this, we masked a penumbra of the cast shadow by black hexagons (Hering, 1874/1964). In addition, since most observers in this experiment were naive to the purpose of the experiment, we provided a clear instruction on the criterion in achromatic setting task.

### 4.1 Instruction for the difference between paper match vs. appearance match

Nine out of twelve observers in this experiment were naive to the purpose of the experiment and had no specialized knowledge about human color vision. We therefore thought that it might be too difficult for them to properly complete the task without careful instructions. Indeed matching criteria have been reported as a major contributor to the degree of color constancy. Arend & Reeves (1986) demonstrated that a performing asymmetric matching based on a paper-match criterion yielded a substantially higher degree of color constancy than matching in terms of color appearance (“hue-and-saturation” match). Though our task did not require a matching per se, the same concern should apply. Also see other studies for focused discussion on this topic (Radonjic et al. 2016; Reeves et al, 2008). Thus to help observers understand the criterion, we set up a few practice trials as shown in Figure 7A. These scenes were shown to each observer one-by-one. The sheet used here contained only achromatic colors, and this sheet was never used in the main experiment. We placed a large white paper at the center of the scene to show how a real white paper would look like when an illuminant color changes. We lit the scene using a single illumination generated by activating only one of the R, G, or B phosphors. Observers were then asked to adjust the chromaticity and the luminance of left and right test fields using different criteria. For the right test field, observers were instructed to adjust the chromaticity and the luminance so that it appears as a full white surface placed under the test illumination (i.e. the matching criterion used in the actual experiment). On the other hand, for the left test field, observers were asked to make a field that simply appears white. Once an experimenter confirmed that the observer understood that it was possible to have two different matching criteria, and that the observer could differentiate two matching criteria, the practice session ended and the main experiment began.

**Figure 7:**
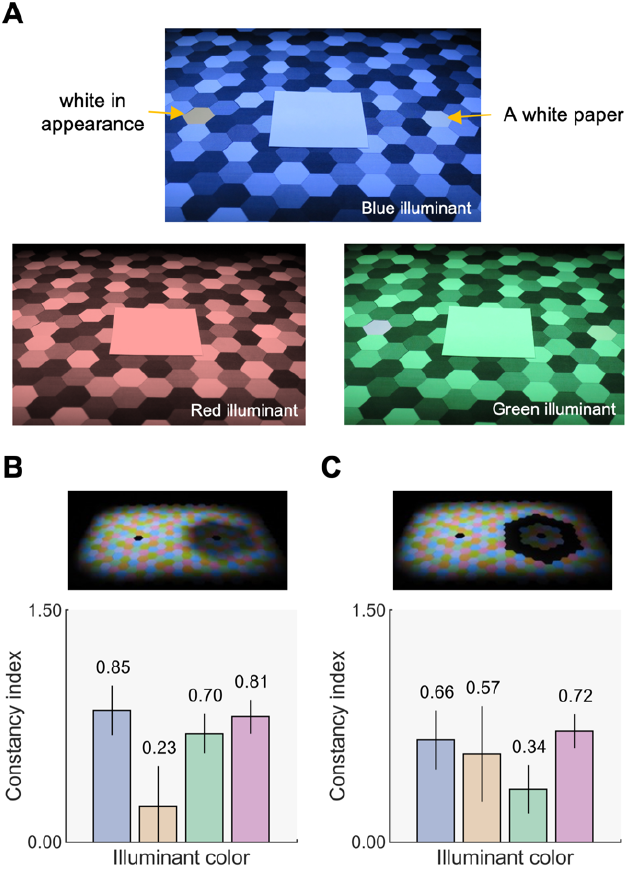
(A) Practice session to help observers to distinguish two matching criteria for achromatic adjustments. (B) Constancy indices for the no-outline condition. (C) Constancy indices for outlined conditions. The bars show mean constancy indices across twelve observers. The error bars show ± 1.0 standard error across twelve observers.

### 4.2 Experimental condition

As shown in Figure 2B, the scene configuration used in Experiment 2 is similar to that of Experiment 1, but we removed a black object at the center of the scene and instead inserted a paper (see Figure 2B, “a paper to create a cast shadow”) which blocked the lights emitted from the left projector to cast a shadow onto the scene. This was because if instead we kept a black object, observers could see that the darkened region is due to the cast shadow and the region would remain perceived as a shadow regardless of our experimental manipulation. As shown in Figure 7B, in the “no-outline” condition, we expected that a shadow would appear as a shadow. In contrast, in an “outlined” condition, we expected that the recognition of shadow weakened as the penumbra of the cast shadow was masked by black hexagons (Figure 7C). One block had 5 skylight color conditions, and one session consisted of two blocks (no-outline, outlined). All observers completed 2 sessions.

### 4.3 Results

Figure 7B and 7C report constancy indices in Experiment 2. To compute constancy indices, because of the limited number of trials for each observer in this experiment, we divided L/(L+M) and S/(L+M) axes by the standard deviation of all observers’ settings for the white skylight condition to approximately equate the scale of both axes. In other words, a common scaling value was used for all observers. We found overall lower constancy indices than those reported in Experiment 1. Again, the influence of skylight color is not apparent. Also, the influence of outlining the cast shadow is not clearly seen. A two way repeated-measures ANOVA was performed with four skylight colors (blue, yellow, green and magenta) and two outline conditions (no-outline and outlined) as the within-subject factors for the *CIs*. The main effects of the color of skylights and outline conditions were not significant (*F*(3, 33) = 1.84, *p* = 0.159, *F*(1, 11) = 0.26, *p* = 0.620, respectively). The interaction between the two factors was not significant either (*F*(3, 33) = 1.34, *p* > 0.278).

These results confirmed that even with a relatively large number of observers color constancy held reasonably well in cast shadows, though the degree of constancy decreased from that in Experiment 1. There was no systematic effect of the recognition of shadow, suggesting that observers were still able to infer the influence of illuminant falling onto the cast shadow even without a strong recognition of the shadow. There was no effect of the skylight color, again showing little evidence for the use of prior knowledge about skylight color.

## 5. General Discussion

It is a mathematically ill-posed problem to correctly judge whether a given material is a bright white material in a blue shadow or a dark blue material under a white illumination. We face this difficulty in everyday life, but there was little experimental work to explore mechanisms that underpin this judgment.

To fill this gap we conducted several psychophysical experiments. Results confirmed that color constancy in shadows holds as good as color constancy in a single-illuminant scene. Furthermore, we found that the experimental manipulations we tried did not affect the degree of color constancy. First of all, the skylight color did not have a strong effect on observer settings. Second, the recognition of a shadow did not affect the degree of color constancy. Each point will be further discussed in subsequent paragraphs.

One interesting observation we found was that the color of skylight had little systematic effect on adjustment results. A computational model that relies on a prior about the typical color of skylight would predict that observer settings are located around the bluish or yellowish region of the color space. Indeed it might be reasonable to expect such a prior because the shadowed region in natural environments presents strong chromatic regularities. In a sunny outdoor environment, the color temperature of the skylight varies only between around 8000K and 25000K, and the variation is tightly restricted along the CIE daylight locus (Morimoto et al., 2022). However, in contrast to such an observation, the observer settings were much better predicted by the color of skylight used in the experiment. This is rather consistent with a controversy around the existence of daylight prior in human color constancy (e.g. Delahunt & Brainard, 2004; Hurlbert, 2019).

In many computer vision applications, color constancy is framed as a problem to find a single appropriate white point in a given scene, which was then used to shift the colors in the whole scene towards a direction to discount the influence of illumination. In some empirical studies, a similar strategy has been suggested as a basis of perceptual constancy (e.g. Helson, 1947; Gilchrist et al. 1999). However, if cast shadows exist in a given scene multiple reference points may need to be locally set. For this reason, a mechanism to compute scene statistics globally (Buchsbaum, 1980; Golz & MacLeod, 2002) would not explain the color constancy we observed in this experiment. Instead, a strategy to segment scenes into small fragments needs to be performed beforehand (Golz, 2008), or such computation might need to be done per object (Hedjar et al., 2023). Local adaptation (von Kries, 1905), as opposed to the global adaptation, would be another mechanism to set multiple anchors across the visual field.

Considering these arguments, a mechanism to detect a cast shadow from the visual field seems to be useful to figure out how a given scene should be segmented into smaller fragments. This is a primary reason why we suspected that the spatial structure of the shadow provides a strong cue for us to find a cast shadow. However, rather surprisingly, outlining the shadow had little impact on color constancy in this study. We predict several reasons for this. Firstly some observers might have still perceived the outlined shadow as a shadow even after violating the spatial intensity regularity. Secondly, the shadowed region might have appeared as being illuminated by a spotlight (Khang & Zaidi, 2004). In either case, observers would be able to hold a separate reference for a cast shadow and other regions in the scene, which would maintain color constancy.

We observed moderately high degree of color constancy in Experiment 2 where mostly naïve observers were recruited, but after the Experiment 2 we suspected that two factors may have played a role in this. One primary factor would be that we gave the explicit instruction regarding adjustment criteria which has been reported in past studies (Reeves, Amano, & Foster, 2008; Radonjic & Brainard, 2016). Secondly, the fact that observers saw that the illuminant color was being changed during the experiment might have also had an effect. To test these ideas, we ran a follow-up experiment that was identical to Experiment 2 except that no practice trials shown in Figure 7A were provided and we asked the observers to leave the experimental room when the illuminant color was changed. Then, we found that constancy indices dropped to nearly zero. Though it is not very clear which factor contribute, or to what extent, as the two factors were manipulated simultaneously; these results showed that instructions and/or experimental procedure have substantial effects on color constancy in shadows, consistent with reports from traditional color constancy studies for single-illuminant environments (Radonjic & Brainard, 2016).

There are some limitations in the study. First, the spatial configuration used in experiments is limited to a set of colored hexagons. We deliberately chose this abstract and simplistic configuration to test our hypothesis in the absence of other cues that might be available in the real world (Granzier et al. 2014). Nevertheless, if instead we used a more complex scene that e.g. contains a wider range of reflectance sets, that might have yielded different results. Indeed the use of realistic environment has been suggested to affect the degree of color constancy. (Granzier et al. 2009; Mizokami, 2019; Radonjic et al. 2015a; Radonjic et al. 2015b). Second, we used a single observer task. As reported in previous studies (Smithson, 2005), the choice of appropriate methodology has been a center of discussion in color constancy literature. While there is no gold-standard method, there might have been an easier task for naive observers. However, we emphasize that because of this difficulty we provided a clear practice session in Experiment 2, which yielded a reasonably good color constancy. Third, there was a naturally limited number of experimental conditions we could test, so we needed to prioritize the present research questions. One additional thing we tested, though not reported in the main text, was whether the use of more saturated hexagons with narrow-band reflectances would affect the degree of color constancy. This additional condition was tested in both of the scene set-ups in Experiments 1 and 2. However, we found that there was no noteworthy difference from the main results and the conclusion stayed the same.

Color constancy has been a core domain in color vision research, and its mechanism has been investigated under many experimental manipulations. However, the real world presents diverse complexity in illuminant conditions, and our understanding of color perception in real-world situations might still be limited in this sense. Future studies are thus expected to reveal mechanisms in the presence of all environmental complexities in which our visual system normally operates. The present investigation of the effect of a cast shadow will potentially contribute to the advancement of our understanding of color constancy.

## Acknowledgement

All authors thank Tanner DeLawyer for editing the language in the manuscript. This work was supported by JSPS KAKENHI Grant Number 19K22881. TM is supported by a Sir Henry Wellcome Postdoctoral Fellowship and a Junior Research Fellowship from Pembroke College, University of Oxford. This research was funded in whole, or in part, by the Wellcome Trust (218657/Z/19/Z). For the purpose of open access, the author has applied a CC BY public copyright license to any Author Accepted Manuscript version arising from this submission.

## Data access

Raw experimental data are available upon a request.

